# Low-frequency ultrasound-mediated blood-brain barrier opening enables non-invasive lipid nanoparticle RNA delivery to glioblastoma

**DOI:** 10.1101/2025.01.23.634427

**Authors:** Maya Elbaz, Nitay Ad-El, Yulia Chulanova, Dor Brier, Meir Goldsmith, Mike Bismuth, Alina Brosque, Divsha Sher, Anna Gutkin, Dana Bar-On, Dinorah Friedman-Morvinski, Dan Peer, Tali Ilovitsh

## Abstract

Ionizable Lipid Nanoparticles (LNPs) are an FDA-approved non-viral RNA delivery system, though their use for brain therapy is restricted by the blood-brain barrier (BBB). Focused ultrasound combined with microbubbles can disrupt the BBB, but delivering large particles requires balancing increased peak negative pressures while maintaining microvascular integrity. Herein, we optimized low-frequency ultrasound parameters to induce high-amplitude microbubble oscillations, enabling the safe delivery of LNPs across the BBB. First, BBB opening was assessed at different frequencies (850, 250, and 80 kHz) and pressures by monitoring the extravasation of Evans blue (~1 kDa). Next, the delivery of 4, 70, and 150 kDa Dextrans, LNPs entrapping Cy5-siRNAs (~70 nm in diameter), and LNPs entrapping mRNA (~100 nm in diameter) was evaluated via microscopy and bioluminescence. In a glioblastoma syngeneic mouse model, siRNA-Cy5-LNP was successfully delivered. A frequency of 850 kHz and 125 kPa pressure induced safe BBB opening, enabling delivery of both small molecules and LNPs. In healthy brains, LNP entrapping siRNAs delivery increased 10-fold compared to controls, and LNPs with mRNAs showed a 12-fold increase in bioluminescence after 24 hours. In glioblastoma tumors, LNPs with siRNAs delivery resulted in a 6.7-fold increase in fluorescence. This study paves the way for non-invasive LNP delivery to the brain, offering a versatile platform for brain therapies.

**TOC graphic:** 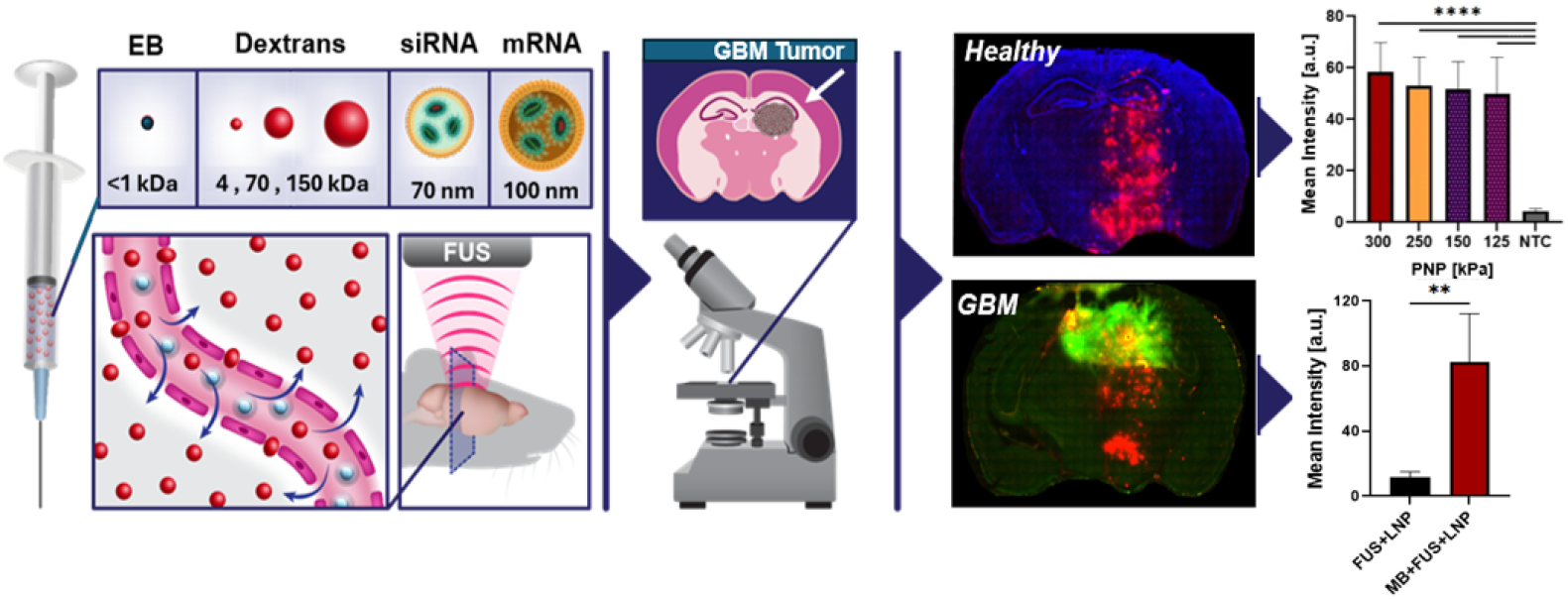

## Introduction

The blood-brain barrier (BBB) is a crucial defense mechanism protecting the brain from pathogens and toxins, yet at the same time poses a significant challenge to the delivery of therapeutics into the brain for the treatment of neurological conditions^1–4^. Glioblastoma (GBM) is the most common and aggressive primary brain malignancy, with a median survival of approximately 15 months following treatments^5^. Although the BBB may be compromised in regions affected by the tumor, this disruption is often irregular and not uniformly distributed across the entire tumor mass^5,6^ Additionally, the extent of BBB compromise is stage-dependent; in the early stages of GBM, the BBB remains relatively intact, preventing the effective passage of large therapeutic agents, including chemotherapies and antibodies^4,7^. Currently, the widely accepted protocol to treat GBM is a combination of invasive surgical resection of the tumor, the chemotherapy temozolomide, and radiation therapy, and it has remained largely unchanged for several years^5^. The high recurrence rates (>90%) and dismal five-year survival rates (<10%) highlight the need for advanced delivery systems capable of effectively transporting therapies across the BBB and into the tumor^8^.

Lipid nanoparticles (LNPs) represent an advanced and FDA-approved form of non-viral delivery system for gene therapy and immunotherapies^9^. In particular, the latest generation of LNPs incorporating ionizable lipids exhibit improved encapsulation and transfection efficiency of both short siRNAs and long mRNAs nucleic acids sequences^10^. Among their advantages are enhanced target specificity, good biodegradability, heightened protein expression, increased therapeutic efficacy and manageable immunogenicity. Consequently, LNPs represent a promising avenue for treating various pathologies through genetic manipulation. Specifically, they are extensively investigated for multiple cancer therapies and are FDA-approved for the indications of Hereditary transthyretin amyloidosis (hATTR) Onpattro^®^ and the mRNA-based COVID-19 vaccines as well as a new RSV Vaccine by Moderna and Pfizer-BioNTech^10,11^. LNPs offer distinct advantages over oligonucleotide-conjugated polymeric particles and liposomes due to their stable encapsulation of genetic payloads within the lipid shell (~close to 100%), safeguarding against off-target interactions and enzymatic degradation, and their superior safety profile and improve production. Additionally, LNPs could outperform adeno-associated viruses (AAVs), which carry the risk of insertional mutagenesis, limitations in sequence size, and constraints in repeated administration^12,13^. Hence, LNPs emerge as promising carriers for gene therapy. Still, their application in advanced brain therapies is impeded by their limited permeability across the BBB, and previous studies utilizing LNPs to treat brain diseases often resort to direct administration into the tumor via stereotaxic injection^14^. Herein, we aim to establish the use of low-frequency focused ultrasound (FUS) combined with microbubbles (MBs) for the noninvasive delivery of LNPs across the BBB and into the tumor.

FUS-mediated BBB opening (BBBO) inherently involves the use of MBs, which are biocompatible ultrasound contrast agents and are FDA-approved for use in humans^15^. The most common MBs are composed of lipid-shell and a gas core with an average radius of 0.75 μm^16^. Upon excitation by FUS, the MBs cavitate, expanding their radius and exerting temporary mechanical forces on adjacent cell membranes. When this process occurs within blood vessels in the brain, the BBB can open safely and transiently, facilitating the local delivery of particles into the brain parenchyma^17,18^. BBBO has been extensively validated through the administration of a diverse array of small molecules for both imaging and therapeutic applications in the brains of rodents and non-primates^19–22^. These studies have consistently demonstrated safe and efficient delivery with minimal off-target effects. Moreover, in recent years, clinical trials are ongoing across a spectrum of neurological conditions, including Alzheimer’s disease, Parkinson’s disease, psychiatric disorders, and chemotherapy/antibody-based treatments in GBM^23–27^.

Despite the significant progress that has been achieved in therapeutic FUS-mediated BBBO, effective delivery of large nanocarriers for advanced brain therapies remains a persistent challenge. Previous efforts to transport genetic payloads using FUS-mediated BBBO, have explored various nanoscale methods, including viral vectors^28,29^, liposomes^30,31^, colloidally stable pDNA-NP^3233^, and even stem-cells^34^. A primary challenge associated with delivering nanoscale particles through the BBB lies in balancing the required peak negative pressure (PNP) to open the barrier while maintaining microvasculature integrity. Higher PNPs are often necessary for delivering large particles, which can lead to undesired microhemorrhages^35^. This is because higher PNPs induces greater MB oscillations, which can create larger gaps in the BBB, facilitating the delivery of larger molecules^36^. Nonetheless, increasing PNP raises the risk of MBs transitioning from stable cavitation, where they oscillate without collapsing, to inertial cavitation, where the bubbles collapse violently, generating microjets that can cause microhemorrhages and brain tissue damage^34,35^. Notably, previous delivery of stem cells (~6 µm) has been reported to induce such transient microhemorrhage^34^. In the context of brain therapies using FUS, low frequency is critically important as it allows higher penetration through the skull non-invasively, combined with lower attenuation passing through tissue, offering an improved performance over ≥1MHz frequencies^37,38^. Our previous research demonstrated that using low frequencies (below 1 MHz, well below the resonance frequency of MBs) enhances the MB vibrational response. By optimizing the parameters, it is possible to select those that allow for the safe BBBO. Previously, we used small molecules such as Evans blue (EB) or gadolinium for extravasation assessment and BBBO confirmation ^39–42^. Now, we aim to optimize parameters for the transfer of LNPs, sized at 70-100 nm with different RNA payloads, using low FUS frequencies. Our goal is to identify the parameter set that allows safe and efficient passage of these particles into the brain following non-invasive systemic injection. BBBO was evaluated across various center frequencies (850, 250, 80 kHz) and PNPs by monitoring the extravasation of EB (~1 kDa). Then, delivery of 4, 70 and 150 kDa Dextrans, LNPs entrapping siRNAs and LNPs entrapping mRNA was assessed using fluorescent microscopy and bioluminescence. Lastly, we demonstrate the successful delivery of LNPs with siRNAs to GBM tumors (Fig. 1).

**Figure 1.**
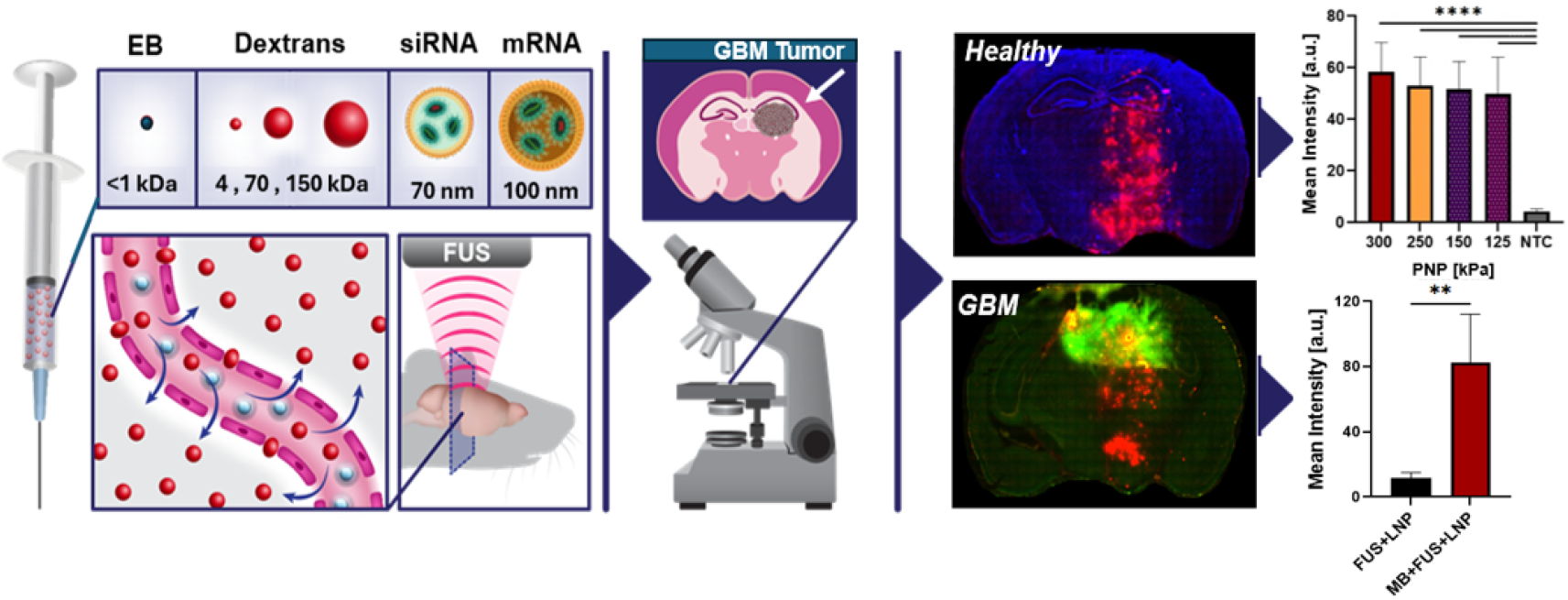
Overview of the non-invasive brain delivery and quantification platform. BBBO was induced using MBs and low-intensity FUS targeted to the right hemisphere. Each time, the mice were systemically injected with one of six particles with progressively increasing size (1, 4, 70, and 150 kDa or 70 and 100 nm) and particles extravasation was quantified using fluorescent microscopy, in healthy brains and in GBM tumors.

## Results

### Focused ultrasound center-frequency optimization

In vivo experiments were conducted using a custom setup, where the mouse was positioned supine, and a laser indicator was used to aid in targeting the right hemisphere (RH) (Fig. 2A). Initial experiments were aimed at determining the optimal center-frequency by monitoring EB extravasation patterns at 850, 250, and 80 kHz in healthy mice brains. For each center-frequency, initial PNP values were chosen based on previous studies or by using the PNP values established at other frequencies as a starting point for testing at lower frequencies. Among the tested frequencies, the 850 kHz center-frequency emerged as optimal for targeted BBBO in the RH, covering an area of ~3.5 × 7 mm^2^ (Fig. 2B,E). The 250 kHz center-frequency resulted in a heterogeneous pattern concentrated only at the brain’s edges (Fig. 2C,F). The 80 kHz center-frequency resulted in a strong, wide focal opening spot, but also caused microhemorrhage at 90 kPa, and only a very mild opening at 75 kPa (Fig. 2D,G, Supplementary Fig.S1). Based on these observations, 850 kHz center-frequency was found optimal in mice brains and selected for further in vivo experiments aimed at the delivery of larger fluorescent particles.

**Figure 2.**
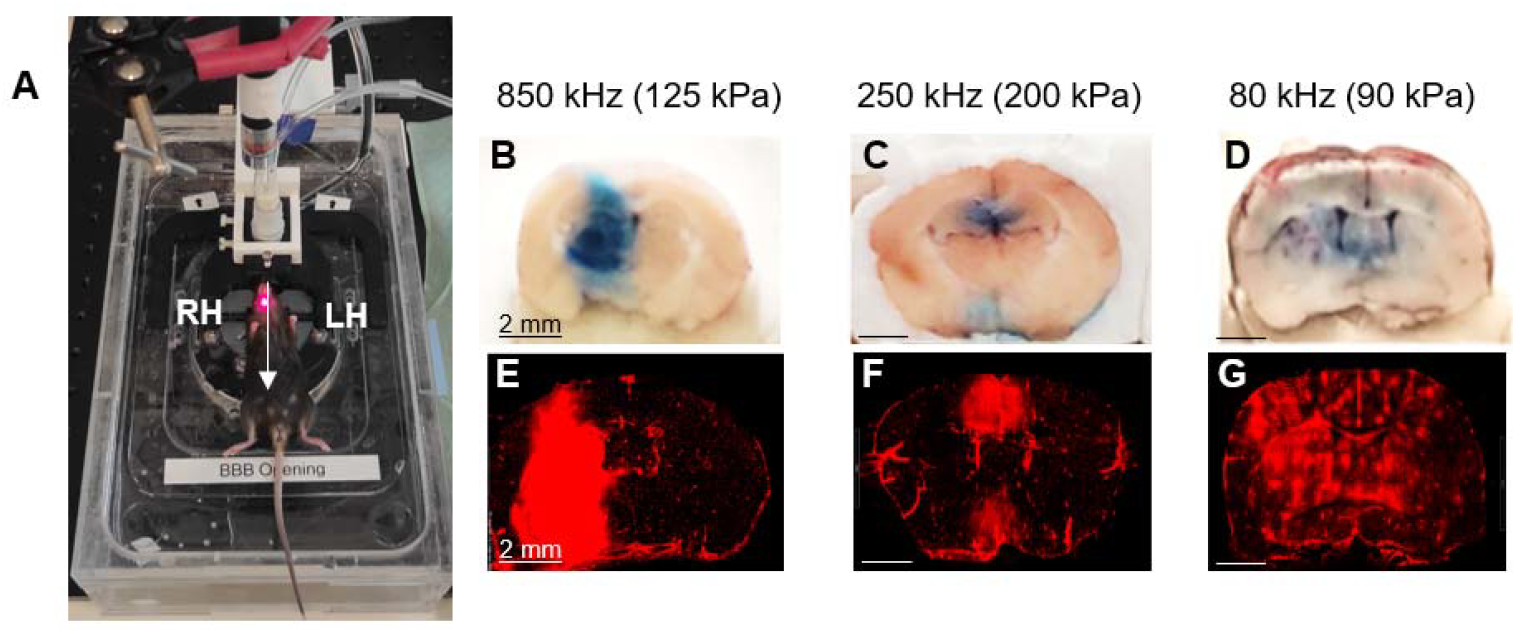
EB extravasation following FUS-mediated BBBO as a function of center-frequency. (A) In-vivo FUS setup that includes a fixed-laser indicator to target the RH. (B)-(G) Evaluation of BBBO at three center-frequencies: Images of EB extravasation in brains treated with a center frequency of (B) 850 kHz, (C), 250 kHz, (D) 80 kHz. (E)-(G) Fluorescence images of the brain slices from (B)-(D). EB extravasation was detected in the red channel. The images were acquired with 20x objective lens. Scale bar: 2 mm

### EB extravasation assessment following FUS-mediated BBBO at 850 kHz

After identifying the center frequency of 850 kHz, optimization experiments for assessing the safe range of PNPs and extravasation assessment were performed. The PNPs were progressively decreased from 500 to 125 kPa until a safe BBBO was achieved, with no microhemorrhage observed in histological analysis. BBBO was visible in the brains’ RH via prone and coronal cuts (Fig. 3A). To compare the size of the BBBO as a function of PNP, the width and height of the opening along the X and Z axes were measured out of the fluorescence images of the coronal brain slices (Fig. 3B, D). Results showed a consistent opening along the Z axis with an average of 6.74 ± 0.13 mm, independent of the applied PNP (not significant), while a significant gradual decrease was observed along the X axis as the PNP decreased from 500 kPa to 125 kPa (5.46 mm ± 0.39 vs 3.49 mm ± 0.29, **** p≤0.0001; One-way ANOVA with Tukey’s multiple comparison). Alternatively, these empirical results can be plotted as an ellipsoid representing the BBBO in each axis as a function of the PNP (Fig. 3C). A lower PNP of 90 kPa was also tested, yet no BBBO was observed (Supplementary Fig.S2). Histological assessment of the presence of blood in each brain section as a function of PNP calculated as percent out of total brain area was used as a measure to evaluate microhemorrhage following treatment, with the amount of blood in control (EB only) sections serving as a reference to healthy brain. Presence of blood in brain histology sections decreased as a function of PNP. At 125 kPa, the values were similar to those of the control, indicating an absence of microhemorrhage and the safety of treatment at this PNP (not significant, One-way ANOVA with Tukey’s multiple comparison) (Fig. 3E-G). Therefore, a PNP of 125 kPa was chosen for the following experiments.

**Figure 3.**
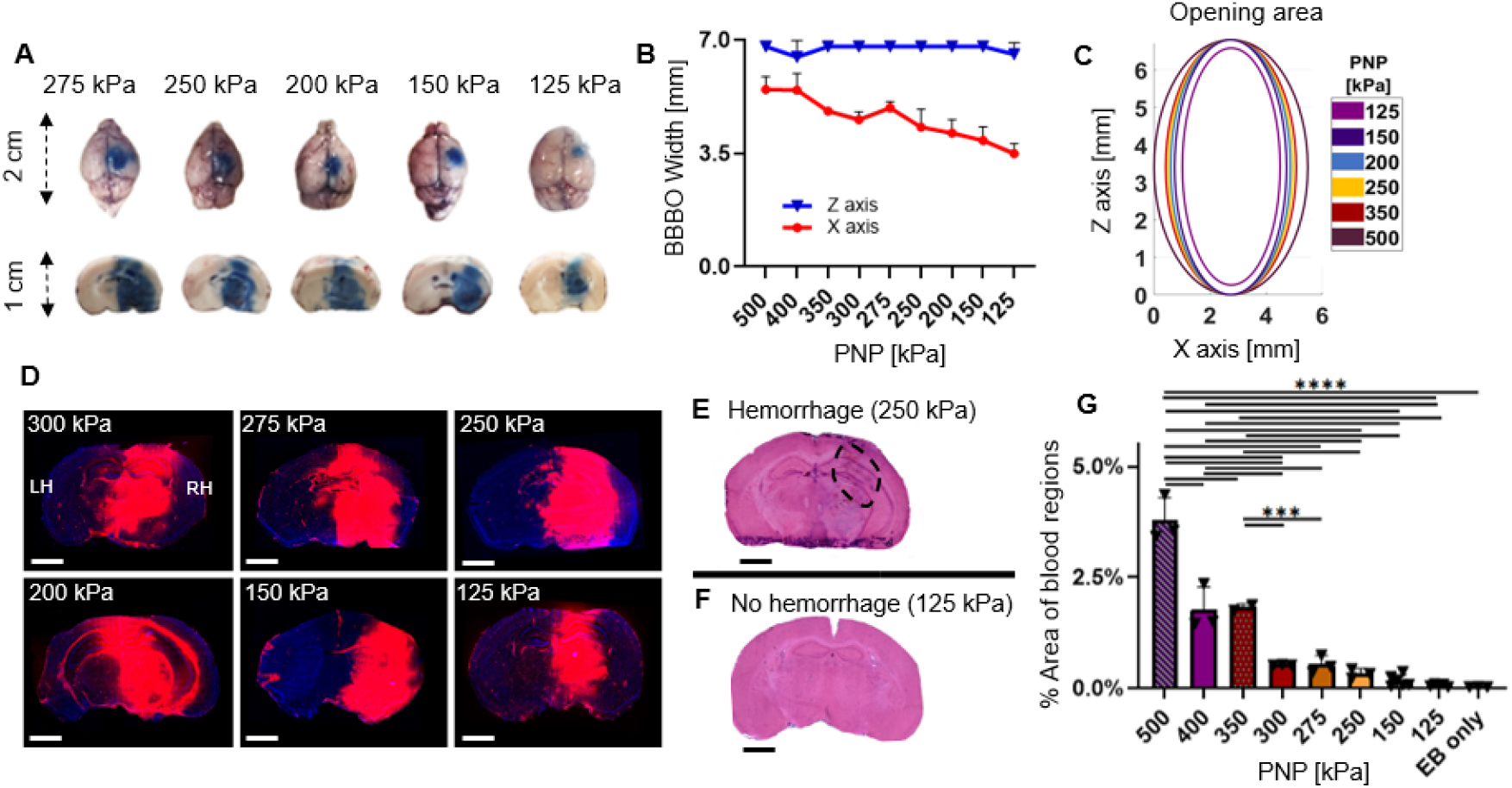
EB extravasation as a function of the PNP at a center-frequency of 850 kHz. (A) EB extravasation in extracted brains positioned prone and in coronal cuts. (B) Comparison of opening width and height [mm] in X and Z axis as a function of PNP. (C) Ellipsoid representation of the opening area, as a function of pressure. (D) EB extravasation imaged using fluorescent microcopy (EB/DAPI Merge). Full brain images were captured in 20x magnification and used to quantify the size of opening. Scale bar: 2 mm. (E-F) Comparison of histological evaluation of microhemorrhages (in H&E staining) at (E) 250 and (F) 125 kPa. (G) Quantification of blood presented in brain slices as a metric to evaluate microhemorrhage area out of the total brain area (%). (G) One-way ANOVA with Tukey’s multiple comparison (*** for p≤0.001, **** for p≤0.0001).

### Fluorescent dextrans brain delivery

After successfully delivering EB (<1 kDa), we used the same FUS parameters to study the delivery of larger dextrans with sizes of 4, 70, and 150 kDa (Fig. 4). The 4 and 70 kDa were Antonia-Red Dextran, which fluoresces in red, while 150 kDa was FITC-Dextran that fluoresces in green (Fig. 4A). Brain slices were imaged using a fluorescent microscope, where the opening size and fluorescence intensity were quantified. Comparison of the opening width along the Z-axis revealed consistent results across all particle types: EB, 4 kDa, 150 kDa and siRNA-Cy5-LNP (not significant), with a small height reduction in the 70 kDa group compared with EB (values were: 6.55 ± 0.36, 6.30 ± 0.31, 6.38 ± 0.33, 6.17 ± 0.35 and 5.6 ± 0.54, respectively). In the X axis, opening with EB (3.49 ± 0.30) was similar to 70 kDa (3.52 ± 0.15), and stronger than 4 and 150 that overlapped (3.05 ± 0.16) (*** for p≤0.001; One-way ANOVA with Tukey’s multiple comparison) (Fig. 4B). Comparative analysis (Two-Way ANOVA with Tukey’s multiple comparison) of fluorescence intensity in the BBBO area compared to control revealed that EB exhibited the strongest intensity among the particles that were tested (Fig. 4C). 4 and 150 kDa dextran presented a similar intensity (not significant), and a stronger intensity over 70 kDa dextran (*** p≤0.001) (with values of: EB: 245.73 ± 6.53 a.u., Dextran 150 kDa: 204.78 ± 28.04 a.u, Dextran 4 kDa: 190.5 ± 19.06 a.u. and Dextran 70 kDa: 133.72 ± 37.1 a.u.). In each group, treated brains exhibited significantly higher marker intensity compared to their control counterparts (**** p≤0.0001) (from left to right: EB: 245.73 ± 6.53 a.u. vs. 13.48 ± 2.38, Dextran 150 kDa: 204.78 ± 28.04 a.u. vs 12.52 ± 4.13, Dextran 4 kDa: 190.5 ± 19.06 a.u. vs 27.42 ± 10.18 and Dextran 70 kDa: 133.72 ± 37.1 a.u. vs 13.65 ± 4.66 a.u.). Notably, the 4 kDa Dextran also showed a diffusion pattern towards the lateral parts of the brain slices, in contrast to the more localized EB distribution represented, however differences were not significant.

**Figure 4.**
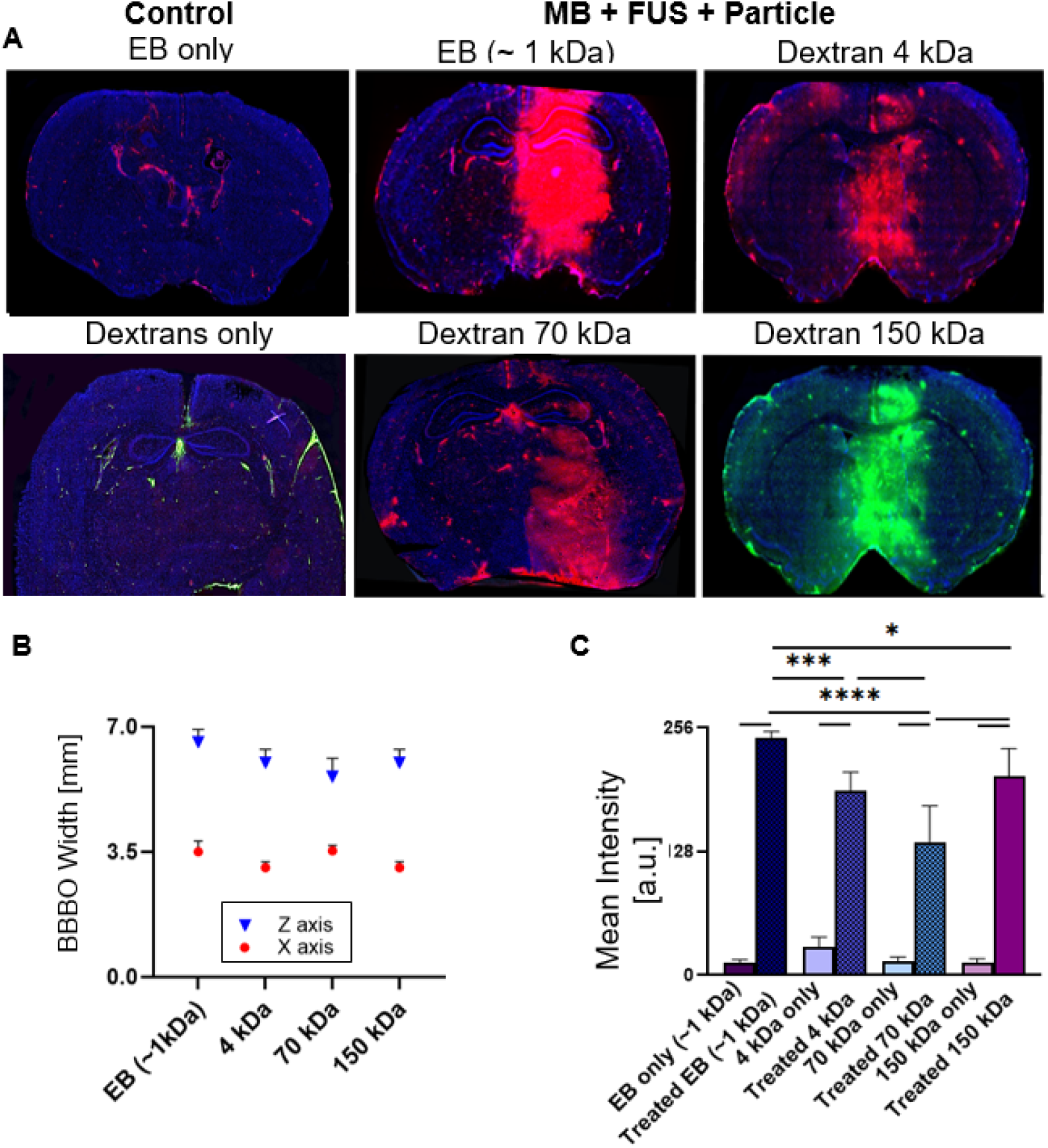
Delivery of EB and 4-150 kDa fluorescent dextran. (A) Fluorescence microcopy images (EB/DAPI Merge) of brain slices following BBBO showing delivery of: left column: controls of EB only, Dextrans only (mix of all three dextrans: 4,70 and 150 kDa), middle and right columns: Treated brains with MB+FUS and: EB (~ 1 kDa), 4 kDa Dextran Antonia Red, 70 kDa Dextran Antonia Red and 150 kDa Dextran FITC. Images were captured in 20x magnification. Scale bar: 2 mm. (B) Size of the opening in the X and Z axes as a function of delivered particle. (C) Fluorescence marker’s intensity comparison between the different molecules and their controls (* for p≤0.05, *** for p≤0.001, **** for p≤ 0.0001.; Two-Way ANOVA with Tukey’s multiple comparison).

### RNA-based LNP brain delivery

The ability to deliver LNPs was then evaluated. The LNPs were constructed using the benchmark ionizable cationic lipid sm-102. siRNA loaded LNPs were produced at an average size of ~70 nm and a zeta potential of −0.426mV, while mRNA loaded LNPs were produced at an average size of ~100 nm and a zeta potential of −0.142mV. Both LNPs had an encapsulation efficiency of above 95% (illustrated in Fig. 5H, Supplementary Fig. S3). The first type of LNP that was used was a non-templet control siRNA-LNP conjugated to Cy5 that was used as a marker to quantify the brain delivery via fluorescent microscopy. The Cy5-siRNA-LNP brain delivery was assessed in the treated group (MB + FUS + LNP) compared to the control group (LNP only) by measuring fluorescence intensity. Since these particles are significantly larger than the dextrans, we started with a higher PNP of 300 kPa and gradually lowered it to the minimum PNP at which a clear opening was achieved without microhemorrhage (Fig. 5A). At the highest PNP of 300 kPa, the opening was accompanied by microhemorrhage, but at 125 kPa, which is the PNP we also used for the dextran delivery, we achieved particle delivery with similar fluorescence intensity to that of the higher PNP, but without histological damage and full recovery of mice (Supplementary Fig. S4). First, a comparative analysis of the mean intensity revealed similar fluorescence signal in the BBBO region for all the PNPs that were tested (not significant), with a significantly increased intensity compared to LNP only control (Fig. 5B) (** p≤0.01, **** p≤0.0001; One-Way ANOVA with Tukey’s multiple comparison). When calculating the opening width in the X axis, measurements were: 4.03 ± 0.42 at 300 kPa, 3.428 ± 0.245 at 150 kPa, and 3.25 ± 0.11 at 125 kPa (Fig. 5C). Importantly, reducing the pressure until 125 kPa did not compromise particles delivery (not significant), and at the same time achieved increased safety with no clinical or histological damage. At the same pressure of 125 kPa, we successfully delivered a variety of dextrans of different sizes to the brain (Fig. 4C, 5D). A direct comparison of all the particles indicates that EB has the highest fluorescence intensity, with the delivery being 18.2 times that of its control. Following that, 150 kDa with a 16.4-fold, siRNA-Cy5-LNP with a 10-fold, 70 kDa with a 9.8-fold and 4 kDa dextran with a 7-fold increase in signal (Fig. 5E, Supplementary Fig. S5) (* p≤0.05; One-Way ANOVA with Tukey’s multiple comparison).

**Figure 5.**
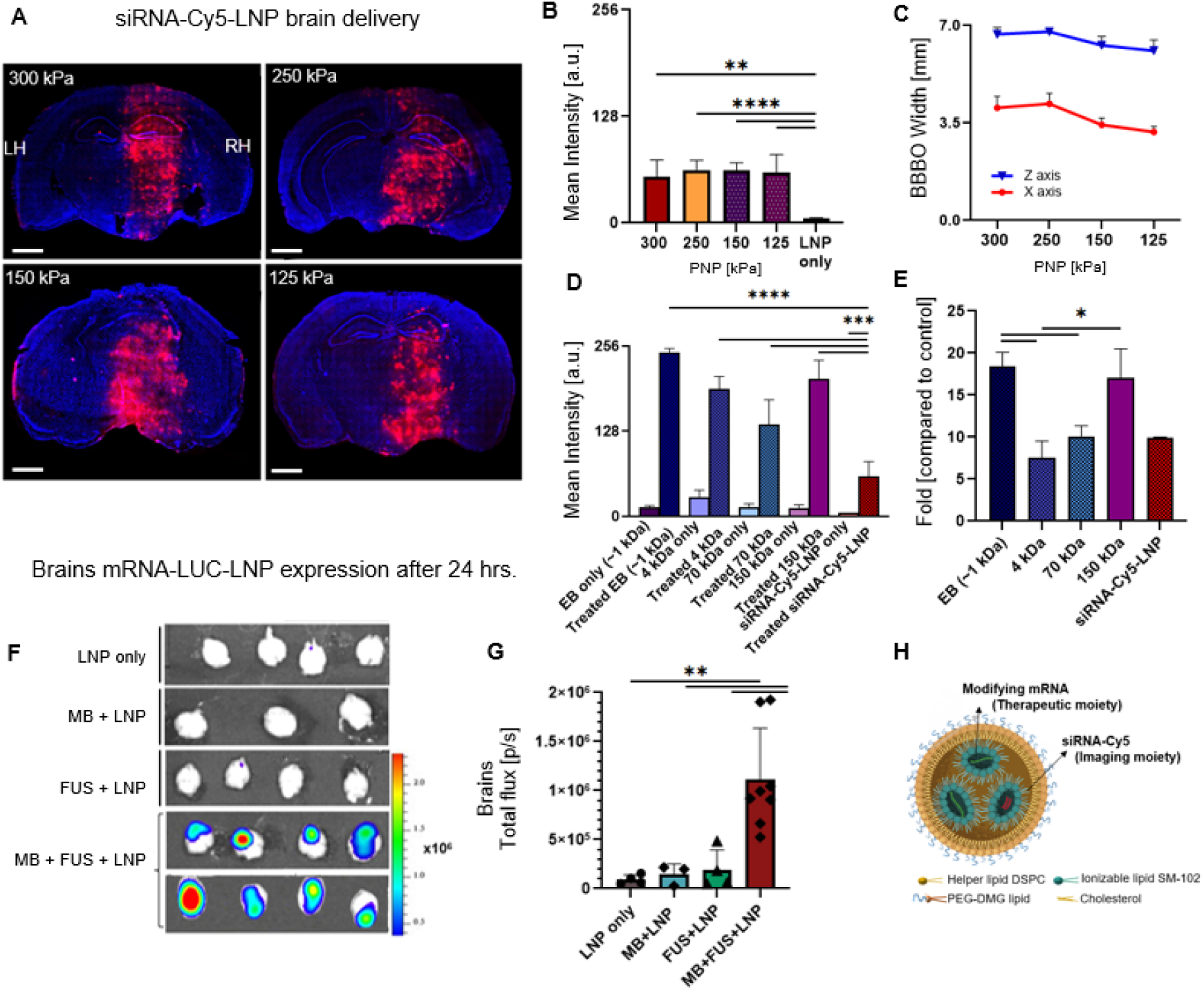
LNPs brain delivery in healthy mice. (A) Fluorescence microcopy images of brain slices after BBBO showing delivery of siRNA-Cy5-LNP (~70 nm) as a function of PNP. Images were captured at a 20x magnification. Scale bar: 2 mm. (B) Comparison of siRNA-Cy5-LNP mean intensity in the BBBO region as a function of the PNP. (C) Size of the opening in the X and Z axes as a function of the PNP. (D) Comparative analysis of all BBBO delivery of 1,4,70,150 kDa particles and siRNA-Cy5-LNP (~70 nm). (E) The graph in (D) presented as the fold increase in intensity compared to the controls. (F) Bioluminescence images of whole brain mRNA-LUC-LNP expression 24 hours post BBBO. (G) Comparison of total flux for the treated group (MB + FUS + LNP) vs controls (FUS + LNP, MB + LNP, and LNP only) in brains. (H) Illustration of the fabricated LNPs (structure and cargo). (D) Two-Way ANOVA with Tukey’s multiple comparison. (B,E,G-) One-way ANOVA with Tukey’s multiple comparison. (* for p≤0.05, ** for p≤0.01, *** for p≤0.001 and **** for p≤0.0001).

After delivering LNPs with siRNAs, LNPs containing mRNA encoding the luciferase protein were fabricated by encapsulation of mRNA-luciferase sequence with SM-102 ionizable lipid based LNPs, with a mean diameter of 100 nm (illustrated in Fig. 5H, Supplementary Fig. S3). These particles facilitate the expression of the luciferase protein and enable the bioluminescent detection of cells that were successfully transfected in the brain. Luciferase expression was evaluated using in vivo imaging system (IVIS) by determining the total flux in whole brains, 24 hours post FUS+MB+LNP treatment (Fig. 5F). The mice group treated with MB + FUS + LNP versus the three control groups (FUS + LNP, MB + LNP, and LNP only) showed a significant increase in the total flux (** p≤0.01; One-way ANOVA with Tukey’s multiple comparison), with a 12-fold increase compared to the LNP-only group (Fig. 5G). Livers were used as positive controls and had no significant difference between the groups (Supplementary Fig. S6).

### Delivery of LNPs into GBM 005 brain tumor

After establishing the capability to deliver LNPs to the brains of healthy mice, the ability to deliver the particles was tested in a GBM mouse model using the same parameters. BBBO was targeted to the RH of GBM bearing mice. The 005 syngeneic GBM mouse model, closely mimics the human disease, and the tumor cells are GFP positive such that they fluoresce in 475 nm range^43^. Fluorescence microscopy confirmed the delivery of siRNA-Cy5-LNP into the tumors (Fig. 6A). Comparison of mean intensity between the treated MBs+ FUS + LNP (n=4) and FUS + LNP (n=3) control groups revealed a 6.7-fold increase in LNP fluorescence signal within the tumor region in treated brains compared to the controls (78.9 ± 28.0 a.u. vs 11.76 ± 3.29 a.u., *** p≤0.001; Unpaired students t-test) (Fig. 6B). Confocal microscopy (x40 magnification) confirmed the co-localization of siRNA-Cy5-LNP signal in the red channel with the 005 GBM cells signal in the green channel (Fig. 6C). The overlap with the blue channel (DAPI) in areas where there is no signal in the green channel suggests that the LNPs were also taken up by other cells in the tumor microenvironment or that there is a decay of the green channel signal as a function of time.

**Figure.**
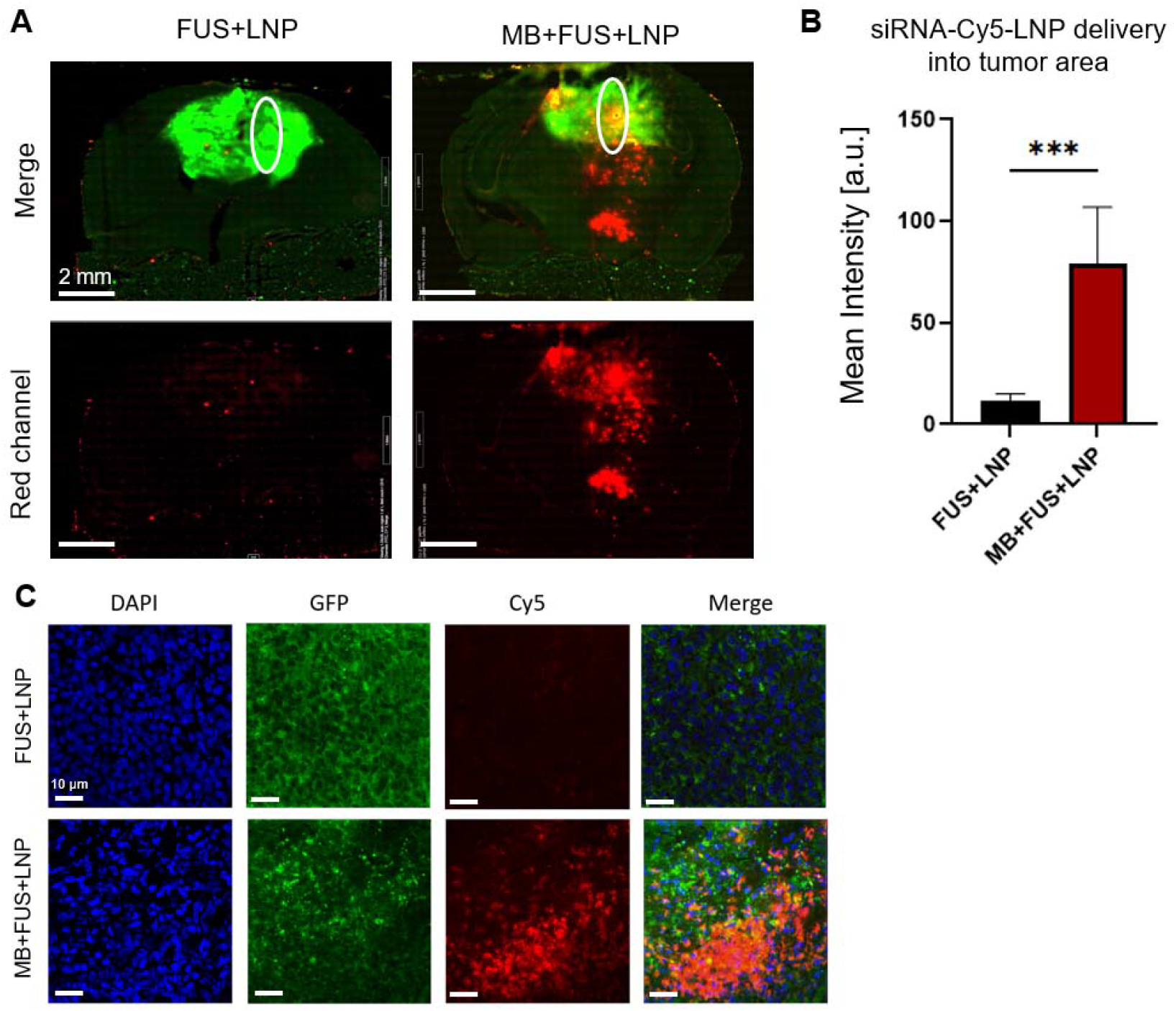
FUS-mediated BBBO for the delivery of siRNA-Cy5-LNP into GBM tumors. (A) Fluorescence microcopy images of brain slices after BBBO showing delivery of siRNA-Cy5-LNP (~70 nm) in treated (FUS+LNP+MB) vs. control (FUS+LNP) GBM brains. Images were captured with a 20x magnification. Scale bar: 2mm. (B) Quantification of siRNA-Cy5-LNP fluorescence in treated compared to controls brains. (C) Confocal microscopy (x40 magnification) of the tumor region stained by DAPI. Green channel shows the GFP positive tumor cells and red channel is the siRNA-Cy5-LNP. Scale bar: 10 µm. (B) Unpaired students t-test (** p≤0.01).

## Discussion

This study investigates the use of low-frequency FUS-mediated BBBO with MBs to enhance the delivery of large particles, focusing specifically on mRNA-based LNP as a platform for future advanced non-invasive local brain therapies. In this work, we developed a robust platform for the noninvasive, safe, and efficient delivery of LNPs with diameters of ~70-100 nm to both healthy brain tissue and GBM tumors. To the best of our knowledge, this is the first demonstration of successful delivery of large LNPs to GBM tumors using FUS-mediated BBBO. This achievement addresses a critical gap in the clinical translation of LNPs, which are FDA-approved and widely recognized as advanced therapeutic agents but have not been used in brain therapies due to the challenges posed by the BBB. While there are 24 clinical trials underway using MB-enhanced ultrasound for gliomas, all focus on small molecules, leaving the potential of LNPs unexplored^44^. Our work demonstrates that LNPs, with their ability to encapsulate RNA-based therapeutics such as siRNA and mRNA, can now be safely delivered to the brain using a noninvasive approach. Additionally, our study presents a comprehensive evaluation of the delivery of a wide range of molecules and macromolecules, including small molecules, dextrans of varying sizes, and LNPs, enabling the identification of a unique set of parameters, that allow for the safe and efficient delivery of both small and large particles across the BBB.

Our results show that operating at a center-frequency of 850 kHz in a PNP of 125 kPa establishes a consistent threshold for safe and effective delivery of particles of various sizes, while preserving tissue integrity. This approach contrasts with prior studies that employed higher frequencies, that are less clinically relevant, resulting in size-dependent delivery, where the need for increased PNPs for larger particles is often limited by safety concerns^34,35,36,45,46^. Moreover, low frequencies offer improved skull penetration efficiency and enable wider focal coverage that can be suited for testing brain delivery. In this study FUS-mediated BBBO with MBs was tested in three low intensity center-frequencies (850, 250 and 80 kHz) by monitoring the extravasation of EB. A MB dose of 2 × 10_ MBs per 20 g of body weight was chosen as it is consistent with our previous studies on BBBO to yield efficient BBBO^40–42^, and falls within the commonly reported range used in BBBO studies in mice and according to the clinically approved dose of FDA-approved commercially available MBs^30,47^. Additionally, the MBs used in this study were size-selected to remove bubbles larger than 5 µm, reducing size heterogeneity and enhancing safety. The use of an identical transducer operating at various frequencies allows for the study of frequency effects without altering other parameters such as aperture size or focal distance. This provides a platform for accurately examining the impact of ultrasound frequency on BBBO. The results presented distinct variation in the opening pattern and coverage area between the different center-frequencies. For each center-frequency, the initial PNP values were chosen based on previous studies or by using the PNP values established at other frequencies as a starting point for testing at lower frequencies. At 80 kHz, we initially tested 180 kPa based on values determined for the 250 kHz center frequency. However, this PNP caused safety concerns, so it was gradually reduced. At 90 kPa, strong BBBO was achieved across the majority of the brain, but microhemorrhages were observed. Further reduction to 75 kPa yielded mild BBBO without microhemorrhages. Based on these results, we concluded that 80 kHz is not an ideal frequency for focused and precise BBBO in the mouse model. At 250 kHz, BBBO was primarily observed at the edges of the brain near the skull (top and bottom), while minimal opening was seen in the center of the brain. This may be due to the phenomenon of standing waves contributing to the effect, suggesting that working with a larger animal model, such as a rat, could mitigate this impact^48^. Moreover, our original study from 2018 identified the safe range of PNPs as below 190 kPa, at which microhemorrhage was observed^39^. When replicating this with a new, identical transducer, we found that a PNP range of 180–200 kPa enabled safe and reproducible BBBO without microhemorrhages, as confirmed by fluorescence microscopy and histological analyses^41^. We believe that this slight variation (less than 15%) is likely due to hydrophone calibration differences and associated uncertainties. At 850 kHz, we started with a PNP of 500 kPa based on observations from studies conducted at a frequency of 1 MHz, where a PNP of was reported to effectively induce BBBO^49^. However, at a frequency of 850 kHz, microhemorrhages were observed at this PNP, prompting a stepwise reduction until a safe and effective BBBO was achieved at 125 kPa. This frequency was found optimal out of the three different frequencies that were tasted, with measured focal dimensions of 3.5 × 7 mm^2^, providing a good coverage of the RH in a mouse brain. As such, this center frequency was chosen for the subsequent sonication experiments.

These optimized parameters (850 kHz, 125 kPa, duty cycle of 0.1%) were then applied for the delivery of various-sized fluorescent dextrans (4, 70, 150 kDa) and two types of LNPs: fluorescent siRNA-Cy5-LNP and bioluminescent mRNA-luc-LNP. Comparative analysis of the fluorescent markers revealed a consistent opening area in X axis, and the strongest fold-increase in mean intensity observed for EB, followed by Dextran 150 kDa, which was the closest to the siRNA-Cy5-LNP (~70 nm). Interestingly, EB is a small molecule (~ 1kDa) known for its high binding to albumin, and 70 kDa dextran has a comparable size to the albumin-EB complex^50^. Our results revealed that EB penetration was much stronger than Dextran 70 kDa, suggesting the involvement of size-independent mechanisms in the delivery process. Along the Z axis, we occasionally observed a phenomenon where the BBBO did not cover the full Z-axis. We think that it may be related to the FUS setup, including factors such as the amount of ultrasound gel applied, or the height of the agar pad. Moreover, it can be related to the angle that the mice head is positioned in the setup.

For Cy5-siRNA-LNP delivery, it has been previously reported that larger particles typically require higher PNPs compared to smaller molecules for effective delivery^35,51^. These studies also indicate the potential for microhemorrhage at such PNPs. As a result, LNP delivery was initially tested at 300 kPa to balance delivery efficiency and safety, yet resulted in microhemorrhage as well. Consequently, the PNP was gradually decreased, and it was found that 125 kPa, the same PNP optimized for EB and other molecules, was sufficient for the safe and effective delivery of LNPs. This demonstrates the effectiveness of our low-frequency ultrasound protocol in enabling efficient delivery across a range of particle sizes while maintaining microvascular integrity. Next, in order to further validate the potential of LNPs as nanocarriers for mRNA encapsulation and gene expression manipulation, the delivery performance of mRNA-LNPs was assessed using bioluminescent mRNA-luc-LNP and subsequent luciferase expression measurement 24 hours post-treatment. The results demonstrated effective delivery exclusively in the FUS-mediated BBBO group with 12-fold delivery enhancement compared to the non-treatment control. Importantly, comparison of the treated group (MB + FUS + LNP) versus the three control groups (FUS + LNP, MB + LNP, and LNP) demonstrated that the enhancement in treatment efficacy is specifically attributable to the combined use of MB and FUS, with neither FUS nor MBs alone producing a significant effect. A limitation of our methodology is the need for brain extraction in order to analyze the delivery signal, therefore, the inability to trace BBB disruption at different time points in whole animals. Developing LNPs decorated with magnetic resonance-based contrast agents could enable the used of magnetic resonance imaging for the monitoring of BBBO. Another limitation is that measuring the fluorescent signal provides a qualitative assessment of the received signal. If further quantification of LNP accumulation in brain tissue is desired, one potential method would be radiolabeling the LNPs, allowing precise measurement of their concentration in tumorous and normal brain tissues using gamma counting or autoradiography^52^. Alternatively, by loading the LNPs with a therapeutic RNA payload, its presence in the brain could be quantified using qRT-PCR or other RNA quantification techniques, such as RNA sequencing, to evaluate delivery efficiency^14,53^.

Based on our optimization results, we assessed the delivery of fluorescent LNPs, targeting their delivery into the RH of the GBM syngeneic 005 mouse model, which closely mimics the human disease^43^. The signal observed in the treated brains showed 6.7-fold increase compared to controls. Ultimately, FUS-mediated BBBO using the optimized parameters allowed the delivery of LNPs into the GBM brain tumors. The focal beam of the transducer used in this study (H-115, Sonic Concepts) covered most of the right hemisphere. By targeting the right hemisphere, we were able to use the left hemisphere as a control, a common methodology in assessments of BBBO^30^. However, within the right hemisphere, the beam covered both the tumor and adjacent healthy brain tissue, meaning that it was not exclusively targeted to the tumor. Instead, our focus was on identifying parameters that enable efficient delivery of LNPs to the tumors. To achieve precise targeting of the tumor alone, a tighter focus would be required. This can be achieved by using a larger aperture transducer, employing a higher frequency, or working with a larger animal model where such a focal spot is clinically relevant. Associated limitations would be that a smaller focal zone would also require beam steering (mechanically or electronically) to ensure complete coverage of the tumor. Additionally, this approach would necessitate image-guided methods, such as magnetic resonance-guided focused ultrasound, to detect the tumor and direct the ultrasound beam accurately. In this study, the tumors were injected stereotactically, and the large focal zone of the beam ensured sufficient spatial alignment with the tumor. An alternative approach to enhance targeting is to develop LNPs with therapeutic content specifically designed to target tumor cells or the tumor microenvironment. However, it may also be possible to design a particle that does not impair performance even when delivered to healthy tissue, making the use of a large focal spot advantageous. This approach could shorten treatment time and ensure complete treatment coverage of the tumor. In this study, our primary goal was to establish a noninvasive, safe, and efficient delivery platform for LNPs to both healthy brain tissue and GBM tumors. Hence, the use of a large focal zone was beneficial for achieving this objective.

Here, we focused on a single BBBO treatment session to assess the delivery capabilities. The feasibility of repeated BBBO treatments depends on the therapeutic agent being delivered. For example, in cases involving one-time treatments such as CRISPR^14^ or AAV^54^, a single BBBO may suffice. However, for therapies requiring periodic administration, such as chemotherapy, repeated BBBO treatments would be necessary. The feasibility of repeated treatments has been demonstrated in previous non-human primates studies^55^, showing that BBBO can be performed multiple times without significant adverse effects. Furthermore, currently there are 24 interventional clinical trials listed on ClinicalTrials.gov, focusing on monthly BBBO treatments accompanying the chemotherapy cycles, with few of them completed showing promising results^44,56^. In future work, testing the safety of our parameters in consecutive cycles, and the impact of repeated treatments of therapeutic RNA-loaded LNPs on tumor progression and therapeutic outcomes is also warranted.

While GBM serves as a representative model in our study, the potential of the method extends far beyond this specific pathology. FUS-mediated BBBO represents a promising avenue for addressing a spectrum of neurological conditions, including Alzheimer’s disease, Parkinson’s disease, psychiatric disorders, and chemotherapy/antibody-based treatment in GBM^23–26^. Traditional approaches to overcoming the BBB often rely on drug mechanisms tailored to specific neurological disorders, limiting their efficacy. Specifically, in brain cancer treatment, local chemotherapy administration into shunted tumors remains a common yet invasive procedure fraught with complications and the development of chemotherapy resistance, highly the need for new therapeutic approaches^57^.

The LNPs delivered in this study are capable of efficiently encapsulate modifying-messenger RNA to manipulate gene expression^14,58^, outperform the efficiency and safety of their alternatives such as adeno-associated viruses (AAVs) and plasmid DNA (pDNA), hence the considerable interest in their use^9^. Their high performance was demonstrated in the development of mRNA vaccines for COVID-19 pandemic^11,59–61^. In addition, they not only enable the delivery of therapeutic mRNA sequences but also facilitates the co-delivery of dye-RNA-conjugated tracers for visualization of extravasation in the brain, potentially enhancing theranostic approaches^62^.

In conclusion, this paper is a comprehensive evaluation of a technology platform for the noninvasive delivery of LNPs to the brain via FUS-mediated BBBO. This work represents a significant step forward in enabling the noninvasive delivery of brain advanced therapeutic platforms. While the therapeutic efficacy of encapsulated RNA payloads in treating GBM tumors is beyond the scope of the current study, it is a key direction for future research. Consequentially, further studies, particularly efficacy and survival studies, are needed. Ultimately, our goal is to translate these promising results into tangible clinical benefits for patients suffering from neurological conditions, thereby addressing a critical unmet need in neurotherapeutics.

## Methods and Materials

### Microbubble preparation

MBs composed of a phospholipid shell and a Perfluorobutane (C4F10) gas core, were prepared using the thin film hydration method as previously reported and briefly summarized here ^39,63^. Two lipids of 1,2-distearoyl-sn-glycero-3-phosphocholine (DSPC; 850365C) and 1,2-distearoyl-sn-glycero-3-phosphoethanolamine-N-[methoxy (polyethylene-glycol)-2000] (ammonium salt) (DSPE-PEG2K; 880129C) (Sigma Aldrich, St Louis, MO, USA) were mixed at a molar ratio of 90:10. A buffer mixture (10% glycerol,10% propylene and 80% saline (pH 7.4) were added to the lipid film and sonicated at 62°C until full transparency. The resulting 2.5 mg/ml MB precursor solution was aliquoted into 1 ml in each vial and saturated with Perfluorobutane gas (C_4_F_10_, Cas no. 355-25-9, F2 Chemicals LTD, UK) to remove air. Upon use, the solution was activated by mechanical shaking with a VialMix shaker (Bristol-Myers Squibb Medical Imaging Inc., MA, USA) and centrifuge to purify MBs with radii smaller than 0.5 µm. Size selection was applied to remove MBs with radii larger than 5 µm. The MBs size and concentration were measured with a particle counter system (AccuSizer® FX-Nano, Particle Sizing Systems, Entegris, MA, USA) and showed median diameter of 1.5 μm and a typical concentration of ~5×10^9^ MB/ml. The MBs were used within three hours of their activation.

### RNA-LNP preparation & characterization

Tracer RNA-LNPs were prepared using a previously described method^64^. Briefly, one volume of lipid mixture (Ionizable lipid SM-102, DSPC, Cholesterol, PEG-DMG at 50:10:38.5:1.5 molar ratio) in ethanol combined with mRNA (1:6 molar ratio RNA to ionizable lipid, either 50% siRNA-Cy5 or mRNA-LUC) in a citrate buffer, pH 4.5 were injected into a NanoAssemblr microfluidic mixing device (Precision Nanosystems Inc., Canada) at a combined flow rate of 12 mL min-1. The resulting LNPs were dialyzed twice using 0.5x phosphate buffered saline (PBS) (Dulbecco’s PBS w/o CA & MG) (pH 7.4) for 16 h and 4h to remove ethanol. Cholesterol, DSPC (1,2-distearoyl-sn-glycero-3-phosphocholine), polyethylene glycol (PEG)–DMG (1,2-dimyristoyl-rac-glycerol) were purchased from Avanti Polar Lipids Inc (Alabaster, AL, USA). Heptadecan-9-yl 8-((2-hydroxyethyl)(6-oxo-6-(undecyloxy)hexyl)amino)octanoate (SM-102) was synthesized in-house as previously described^64^. mRNA sequences were purchased from TriLink (San Diego, CA, USA), and synthesized with complete N1-methyl-pseudouridine nucleotide substitution. siRNA-Cy5 was purchased from Integrated DNA Technologies, Inc. The resulting LNP sizes were characterized by dynamic light scattering (DLS) (Zetasizer Ultra, Malvern, Panalytical, Westborough, MA, UK). Zeta potential was determined using the Zetasizer S system (Malvern, Worcestershire, UK). Encapsulation efficiency was determined using the RiboGreen essay (Thermo Fisher Scientific, Waltham, MA, USA). Concentration and size distribution were performed on diluted samples (1:5,000 in PBS) using NanoSight (NTA, NS300, Malvern, UK).

### FUS setup

The FUS setup was composed of spherically focused single-element transducer (H115, Sonic Concepts, Bothell, WA, USA) operated by a transducer power output system (TPO-200, Sonic Concepts) that was located at the bottom of a water tank facing upwards^40,41^. The transducer had a diameter of 64 mm and a focal distance of 45 mm. It supported center frequencies of 80, 250, and 850 kHz via custom matching networks (purchased from Sonic Concept). The PNPs for each center frequency were calibrated using a needle hydrophone (NH0500, Precision Acoustics, UK). For the in vivo experiments, the mice were positioned on top of an agarose spacer. This spacer was prepared by dissolving agarose powder (A10752, Alfa Aesar, MA, USA) in distilled water to achieve a 1.5% concentration, followed by heating to completely dissolve the agar powder and remove gas bubbles. The solution was poured into a mold of 5 cm x 5 cm x 1 cm (length x width x height) and cooled at room temperature. The spacer was placed on top of the water tank as a mouse bed. To precisely target the RH, the setup was equipped with a vertically fixed laser pointer to co-align with the transducer’s focal spot (Fig. 1A).

### In-Vivo BBBO Experiments

Eight to twelve-week-old female C57BL/j6 mice, weighing between 19-23 grams (Harlan, Jerusalem, Israel), were used for the in-vivo FUS-mediated BBBO experiments. All animal procedures were approved by the Institutional Animal Ethics Committee at Tel Aviv University (approval number TAU-MD-IL-2401-103-5) and carried out in accordance with established guidelines.

#### Animal preparation

Mice preparation for the BBBO procedure was as following: all mice were anesthetized with 2% isoflurane using a low-flow vaporizer system (120 ml/min, SomnoFlo, Kent Scientific, Connecticut, USA). Their heads were fully shaved with a machine, and any remaining hair was removed using hair removal cream (Veet, Reckitt Benckiser, France), that was applied for 40 seconds and then removed with a water-soaked pad. The region of interest in the RH was marked by a dot with a marker to assist in the positioning of the mice.

#### BBBO

For the BBBO procedure, mice were systemically injected with 2 × 10^7^ MBs per 20 g of body weight, in 50 µl of degassed PBS (Dulbecco’s PBS w/o CA & MG). US gel was applied on top of the agarose spacer, and the mouse’s head was positioned supine on top of the gel. The FUS treatment was operated within 60 seconds from MBs injection in varied parameters depending on the tested center-frequency. FUS treatments in 850 kHz center-frequency comprised of 1 ms bursts and RPF of 1 Hz (duty cycle of 0.1%), testing nine PNPs: 500, 400, 350, 300, 275, 250, 200, 150, 125 and 90 kPa (n=18 in total). In 80 kHz center-frequency, 3.25 ms bursts and RPF of 1 Hz were operated (duty cycle of 0.1%) and the tested PNPs were: 180, 120, 90, 75 (n=8). The PNPs were optimized for 850 kHz and 80 kHz, by gradually decreasing the PNP until safe BBBO was observed, without signs of microhemorrhage in histology. For 250 kHz, the PNP was set at 200 kPa with the same PRF (1Hz) based on previous optimization^40^. Immediately after the FUS treatment, mice were systemically injected with one out of six different molecules as described next.

#### 1. EB delivery

EB (E2129, Sigma Aldrich) dye solution of 2% in PBS at 4 ml/kg and was systemically injected and allowed to circulate for a duration of 28 minutes before mice were sacrificed. This time point was selected based on previous research^40^. Once identified, the optimized BBBO parameters (850 kHz, 125 kPa, 1 ms bursts, duty cycle of 0.1%) all subsequent experiments were conducted using this protocol.

#### 2. Dextrans delivery

For experiments with dextran molecules, three distinct molecular weights (4 kDa, 70 kDa, and 150 kDa), each labeled with a fluorescent moiety were used (Antonia Red-lysine-dextran [ARLD4, ARLD70], TdB Labs AB, Uppsala, Sweden, and 150 kDa FITC-Dextran [68042-46-8], Sigma-Aldrich). To maintain consistency, each dextran dose was 1 mg in 100µl and circulation time was 10 minutes before scarification^35,65,66^. Mice were divided into three groups (n=5 in each): treated group injected with 1 mg of red-labeled 70 kDa dextran; a second treated group injected with a mixture of red-labeled 4 kDa and green-labeled 150 kDa dextrans, and a control group injected with a mixture of all three dextrans.

#### 3. LNPs delivery

For experiments with LNPs, two types of LNPs were fabricated composing of either 50% siRNA-Cy5 or mRNA-LUC. For siRNA-Cy5-LNP, a fixed dose of 1mg/kg was systemically injected, and brains were harvested 2.5 hours post-treatment. A total of 15 (n=3 each) mice were divided into four groups treated at descending pressures (300, 250, 150, 125 kPa) and NTC. In the case of mRNA-LUC-LNP, a fixed dose of 1mg/kg dose was systemically injected, and mice were imaged 24 hours post BBBO. A total of 21 mice, all injected with mRNA-LUC, were divided into four groups: LNPs only (n=6), MBs + LNPs (n=3), FUS + LNPs (n=4), and MBs + FUS (125-135 kPa) + LNPs (n=8). 24 hours post treatment, prior to the IVIS imaging the mice were injected intraperitoneally with XenoLight d-luciferin (15 mg/kg) (122799, PerkinElmer Inc.)^14^. Post-sacrifice, brains and livers were extracted and imaged for LUC signal. The bioluminescence analysis was preformed using the Living Image software comparing the samples total flux [p/s].

At the end point of each experiment) EB / Dextrans / LNPs), the brains were collected and positioned on top of a Tragacanth Gum paste, which had been prepared by mixing Tragacanth Gum powder (G1128-100G, Sigma-Aldrich) in distilled water at a concentration of 15% (w/v). Subsequently, the samples were flashed-frozen in 2-methylbutane (Sigma-Aldrich) using liquid nitrogen and stored in a −80ºC refrigerator until cryo-sectioning to 20 µm slices.

### GBM 005 orthotopic model

Mouse-derived glioma cell line (GBM 005, GFP+, LUC+), a gift from Prof. Dinorah Friedman-Morvinski, were established using lentiviral transduction of H-Ras and activated Akt in Cre-GFAP/p53+/− C57BL/6 mice, as previously described^43^. Maintained in stem cell medium, specifically DMEM/F:12 medium supplemented with 1% Glutamax (100X), 1% penicillin-streptomycin, B27 supplement (Invitrogen), N2 supplement (Invitrogen), heparin (50 μg/mL), EGF (20 ng/mL), and FGF2 (20 ng/mL), the GBM 005 cells were cultured as spheres and split every 3-4 days using TrypLE Express dissociation reagent (Gibco Corp, 12604-013, Grand Island, NY, USA) when reaching 90% confluency. A total of 30 eight-week-old female C57BL/6JOlaHsd mice (Envigo, Jerusalem, Israel) were anesthetized using isoflurane, positioned in the Kopf Stereotaxic Alignment System, and inoculated with 3×10^5^ GBM 005-GFP-luciferase cells in a 1.5-µl volume using automatic syringe pump in a rate of 0.3 µl/min. Injections were made to the right frontal lobe: ~1.5 mm lateral, 2 mm caudal from bregma, and at a depth of 2.3 mm. Tumor inoculation and growth monitoring was performed by bioluminescence imaging (IVIS Spectrum, PerkinElmer Inc.) every 5 days post tumor cell implantation until the experiment between days 17-21. XenoLight d-luciferin was injected at 15 mg/kg intraperitoneally and mice were imaged within 10 to 30 mins post injection. Bioluminescence analysis was conducted using the Living Image software (PerkinElmer Inc.) comparing the subject’s total flux^14^.

### BBBO for siRNA-Cy5-LNP delivery in GBM 005 orthotopic model

The BBBO experiment in GBM mouse model was performed between days 17-21 post tumor inoculation. Experiment followed the same protocol described for siRNA-Cy5-LNP delivery in healthy mice: The optimized BBBO parameters were 850 kHz, 125 kPa, 1 ms bursts, duty cycle of 0.1%; siRNA-Cy5-LNP was systemically injected at a fixed dose of 1mg/kg, and brains were harvested 2.5 hours post-treatment. Fluorescent microscopy was used to confirm the LNPs accumulation within the GBM 005 +GFP positive tumor region (detailed under ‘Microscopy imaging and Quantitative analysis’). Comparative analysis of siRNA-Cy5-LNPs extravasation in the tumor area in treated vs control brains was conducted.

### Microscopy imaging and Quantitative analysis

Frozen brains were cryo-sectioned to 20-μm-thick coronal in a −20º C cryostat microtome (CM1950, Leica Biosystems). The sections were placed on standard microscope slides and kept in a slide box at −20º C until use. Upon imaging, the brain slides were thawed to room temperature and imaged within 1 hour to avoid dye diffusion. All full brain images in this study were obtained using a hybrid automated microscope (Revolution, Echo, San Diego, USA). The imaging process involved stitching 20 × 30 tiles, each measuring 0.432 mm x 0.36 mm, to create a full slice scan. These scans were conducted at 20x optical magnification. For fluorescent images the following excitation wavelengths and exposure times were used: DAPI (365 nm, 90 ms), GFP (410 nm, 460 ms), Cy5 (690 nm, 790 ms), and Evans Blue (690 nm, 90 ms). Confocal microscopy images were acquired with confocal microscope (IX-83, Olympus) using UPLXAPO x40 objective (NA 0.95) in z-stacks (1 um each, 8 stacks). The nucleus was labeled with DAPI (405 nm), GBM 005 were +GFP positive (488 nm), siRNA-LNP were labeled with Cy5 (640 nm), and subsequently merged using ImageJ® software (Fig. 5C).

Measurement of the width of the opening and the full brain distances in the X-axis and the Z-axis were conducted using the microscope software (Figs. 2B, 3B and 4D). Segmentation of the opening area with EB as a function of pressure in the representation of an ellipsoid was calculated using an ellipse function in MATLAB (version 2018a, MathWorks, Natick, MA, USA) (Fig. 2F). The values were normalized to average mouse brain size. Quantification of opening size area, microhemorrhages and markers intensity were conducted using ImageJ^®^ software (National Institutes of Health, Bethesda, MD). Full-brain fluorescence microscopy images were first imported into ImageJ. The area of opening in red or green channel and the total brain area were manually selected. The opening area was then calculated as a percentage of the total brain area (Figs. 2G, 3C, and 4A). Markers’ intensity was calculated by selecting an opening area at the same size across all brain images (Fig. 3D, 4C). Quantification of BBBO-induced hemorrhage was conducted on full-brain brightfield microscopy images (without H&E staining) using the IHC toolbox in ImageJ. To reduce background noise, each hemorrhage center was defined as 200 pixels^2^, and the IHC toolbox was trained to recognize relevant shades of brown from total brain area (Fig. 2H, S2).

### Histology

Hematoxylin & Eosin staining; The safety of the BBBO treatments was assessed by a standard Hematoxylin (Leica 3801542) and Eosin (Leica 3801602) (H&E) staining of the 20-μm-thick frozen brain sections and scan x20 using the brightfield channel (Fig. 2D-E). DAPI Staining; Tissue slices were mounted onto glass slides and coverslipped after the application of three drops of Mounting Medium with DAPI (Fluoroshield, ab104139, Abcam). The slides were allowed to develop for 15 minutes before further analysis.

## Statistical analysis

Prism 10.1.2 (GraphPad Software) was employed for the statistical analysis. A two-sided Student’s t-test was utilized to compare two experimental groups. In experiments involving multiple groups, differences among multiple populations and sub-populations were assessed using One-Way and Two-Way ANOVA with Tukey’s multiple comparisons. A value of p≤0.05 was considered statistically significant. Differences are presented on graphs in the following abbreviations: blank. for not significant, * for p≤ 0.05, ** for p≤0.01, *** for p≤0.001, and **** for p≤0.0001.

## Author Contributions

Conceptualization: ME, DP, TI; Data curation: ME, NAE, YC, DB, MG, MB, AB, DS, AG; Funding acquisition: DBO, DP, TI; Methodology: ME, NAE, YC, MG, DBO, DFM, DP, TI; Investigation: ME, NAE, YC, MG, DP, TI; Resources: AB, DS, DFM; Visualization: ME, YC, MG; Supervision: DP, TI; Writing—original draft: ME, YC,MG, DP, TI; Writing—review & editing: ME, YC,MG, DFM, DP, TI

## Data Availability

The datasets generated during and/or analyzed during the current study will be made available upon request.

