## Supplementary Figures for "Low-frequency ultrasound-mediated blood-brain barrier opening enables non-invasive lipid nanoparticle RNA delivery to glioblastoma"

**This PDF file includes:**
Supplementary Figures
Fig. S1-S6

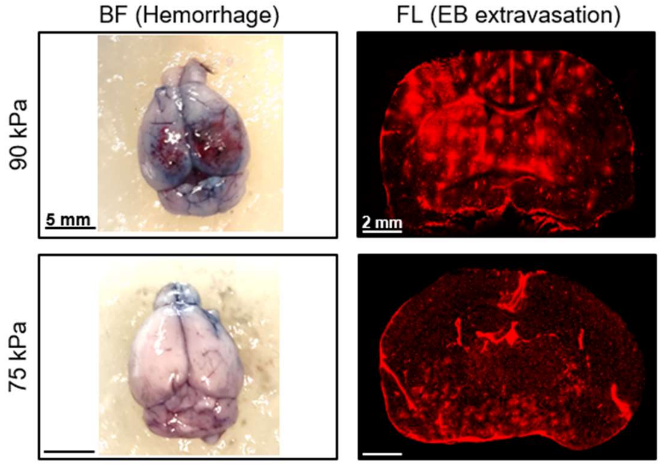


**Figure S1. BBBO at a center frequency of 80 kHz.** Representative images of hemorrhage in BF at 90 kPa and respective EB signal in fluorescent microscopy (in the Red channel), compared to mild extravasation in 75 kPa without a damage in BF. Scale bar: 2 mm (magnification: x20).

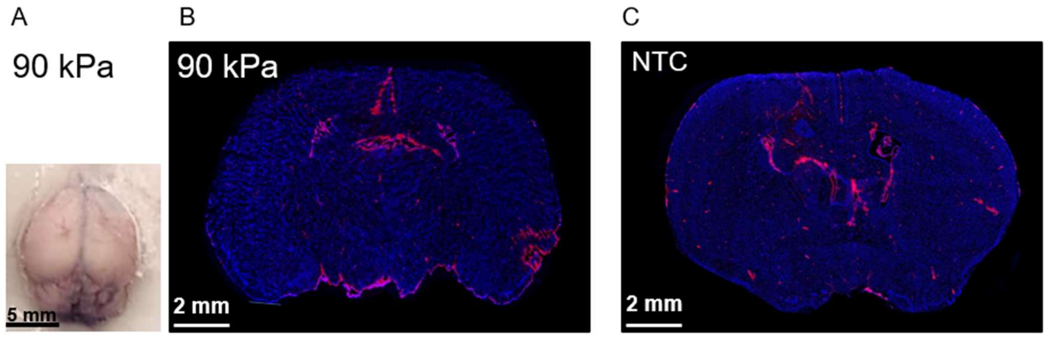


**Figure S2. No BBBO was detected when using an 850 kHz center-frequency and a PNP of 90 kPa.** (A) Top view of an extracted brain. Fluorescence microscopy images of brains treated with: (B) 90 kPa and center frequency of 850 kHz, and (C) Non-treated control (NTC) brain (injected with EB only). All microscopy images were stained with DAPI and acquired at x20 magnification. Scale bar: 2 mm.

siRNA-Cy5-LNPs

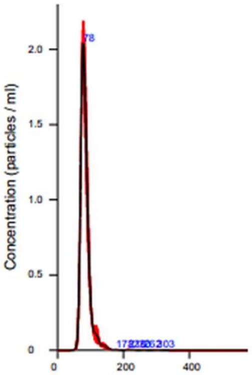


| | |
|---|---|
| Mean: | 84.0 +/- 0.2 nm |
| Mode: | 77.9 +/- 1.1 nm |
| SD: | 17.3 +/- 1.2 nm |
| D10: | 69.6 +/- 0.5 nm |
| D50: | 80.6 +/- 0.6 nm |
| D90: | 100.5 +/- 2.4 nm |
| Concentration (Upgrade): | 5.03e+08 +/- 4.99e+07 particles/ml |

mRNA-LUC-LNPs

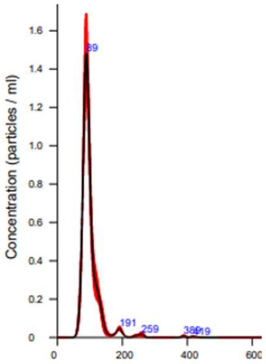


| | |
|---|---|
| Mean: | 100.5 nm |
| Mode: | 88.9 nm |
| SD: | 34.7 nm |
| D10: | 77.2 nm |
| D50: | 92.5 nm |
| D90: | 127.1 nm |
| Concentration (Upgrade): | 4.80e+08 +/- 1.29e+07 particles/ml |

siRNA-Cy5-LNPs:

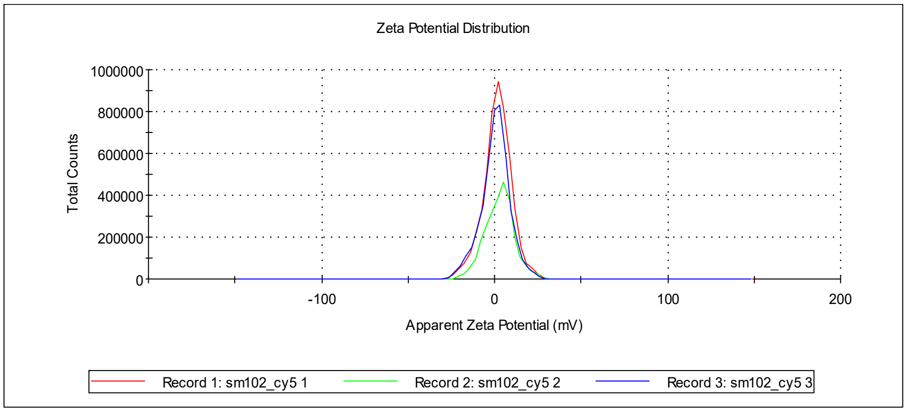


mRNA-LUC-LNPs:

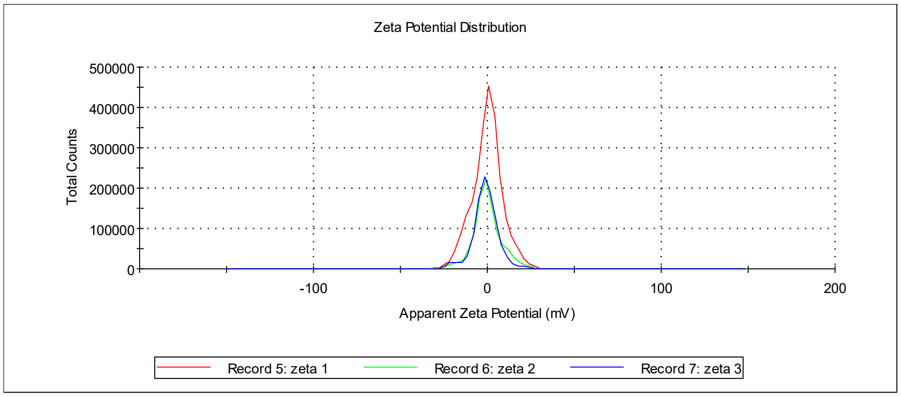


**Figure S3.** Representative measurements of size distribution and concentration of siRNA-Cy5-LNPs and mRNA-LUC-LNPs using NanoSight (samples are diluted 1:5,000 in PBS) and Zeta potential using Zetasizer.

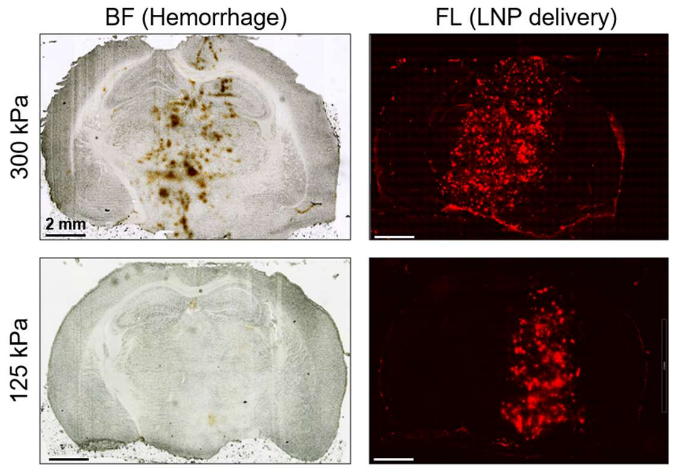


**Figure S4. BBBO at a center frequency of 850 kHz.** Representative images of hemorrhage in BF at 300 kPa and respective siRNA-Cy5-LNPs signal in fluorescent microscopy (in the Red channel), compared to reduced delivery of the LNPs in 125 kPa without a damage in BF. Scale bar: 2 mm (magnification: x20).

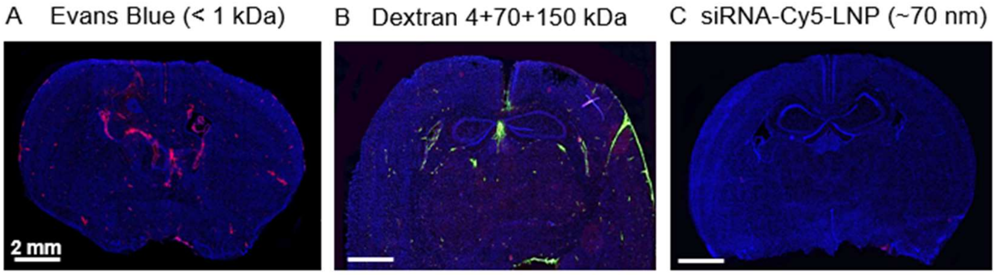


**Figure S5.** Representative fluorescent images of non-treated control brains: (A) with Evans blue, (B) with a mixture of dextran 4 kDa (Antonia-Red), 70 kDa (Antonia-Red) and 150 kDa (FITC), and with (C) siRNA-Cy5-LNP (~70 nm). All images were stained with DAPI and acquired at x20 magnification. Scale bar: 2 mm.

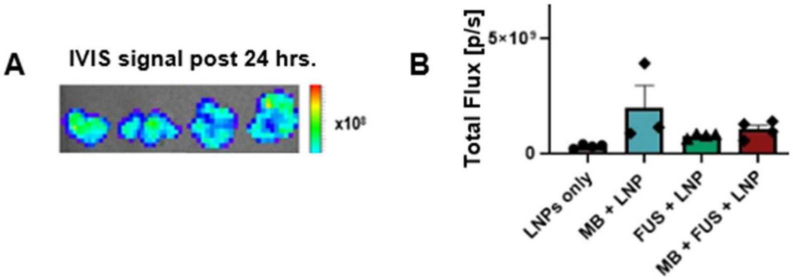


**Figure S6.** Confirmation of mRNA-sm102-LUC-LNP injection by positive liver controls. (A) Representative IVIS image of mRNA-sm102-LUC-LNP uptake into liver in the treated group (MB+FUS+LNP). (B) Comparison analysis of mRNA-sm102-LUC-LNP uptake to the livers (total flux [p/s]) between the different groups (ns).
